# Daily transient coating of the intestine leads to weight loss and improved glucose tolerance

**DOI:** 10.1101/2021.06.11.447961

**Authors:** Tammy Lo, Yuhan Lee, Chung-Yi Tseng, Yangshuo Hu, Christos S. Mantzoros, Jeffrey M. Karp, Ali Tavakkoli

**Author notes:** Corresponding Authors: Ali Tavakkoli, M.D., Brigham and Women’s Hospital, 75 Francis St., Boston, MA 02115, United States, Jeffrey M. Karp, Ph.D., Brigham and Women’s Hospital, 60 Fenwood Rd., Boston, MA 02115, United States. Authors equally contributed (Tammy Lo, Yuhan Lee).

## Abstract

The worsening obesity epidemic has driven a parallel increase in incidence of type-2 diabetes (T2D) found in 13% of the adult US population. Roux-en-Y gastric bypass surgery (RYGB) is the golden standard surgical treatment for obesity-related T2D. However, over 70% of T2D patient do not meet the NIH criteria for surgery, and many that do, are reluctant to consider such an invasive intervention. Mechanisms behind the antidiabetic effect of RYGB are not clearly understood, but isolation of proximal bowel from nutrient exposure plays a critical role. In this study, we tested the impact of daily administration of an oral gut coating agent, LuCI (***Lu***minal ***C***oating of the ***I***ntestine), in controlling weight and insulin resistance. LuCI is not systemically absorbed and provides a transient coating of the proximal intestines and excludes the coated bowel from nutrient contact, thus mimicking the effects of RYGB. We demonstrate that LuCI results in weight loss and improved insulin sensitivity in a rat DIO model. LuCI also appears to replicate hormonal changes seen after RYGB, by increasing post-prandial GLP-1 and GIP levels. LuCI has the potential to replicate the metabolic benefit of surgery and be a novel therapeutic strategy for obesity associated T2D.

## Introduction

The modern epidemic of obesity and its myriad of associated comorbidities have raised global health concerns. Most notably, obesity significantly increases the risk of developing type 2 diabetes (T2D) by 40 fold for obese women with body mass index (BMI) > 30, and by 80 fold for BMI > 35 ^1,2^, affecting over 650 million people worldwide, leading to substantial mortality, morbidity and vast healthcare expenditure^3^. First-line therapy for T2D relies on oral antidiabetic drugs; however, disease progression to dependence on insulin injections is common.^23^ Although anti-diabetic medications are available, despite decades of drug refinement for T2D, a recent study shows that over the 10 years following diagnosis, T2D patients spent one-third to over one-half of their time in suboptimal glycemic control and that reducing this time could lower the risk of microvascular and macrovascular complications.

Recently, bariatric surgery and specially Roux-en-Y gastric bypass (RYGB) has been shown in multiple randomized clinical trials to be superior to medical treatment or lifestyle intervention in the management of T2D^4– 7^. Despite these overwhelming evidence only a small percentage of T2D patients however undergo RYGB, as many do not meet the current NIH guidelines for surgery while others are concerned about short and long term complications following bariatric surgery^8^.

There has therefore been significant interests in developing less invasive alternatives, that can replicate the metabolic success of RYGB^9^. Following RYGB, it has been reported that over 80% of patients achieve complete T2D remission. Furthermore, studies in the field suggested that the impact of RYGB on T2D cannot be solely justified by the effects of weight loss and restrictive energy intake alone^10^. Weight-independent anti-diabetic effects brought on by RYGB are apparent following the very rapid resolution of T2D^11^. Various underlying mechanisms have been suggested, including the regulation of incretin hormones^12^, secretion of bile acids^13^, enhancement in intestinal nutrient sensing^14,15^, and alterations demonstrated in the gut microbiome^16,17^. These well documented mechanisms not only involve hormonal and physiological adaptations following RYGB^13,15,18,19^, but the acknowledgement that isolation of the proximal bowel from nutrient exposure is also a critical part of surgical success^20,21^. This led to our approach to develop a novel and orally administered, gut barrier coating that seeks to reproduce the impact of RYGB without the need for an invasive procedure and its associated risks.

We have developed LuCI (in short for **<u>Lu</u>**minal **<u>C</u>**oating of the **<u>I</u>**ntestine), a candidate therapeutic for diabetes that transiently coats the gastrointestinal mucosa, independent of the pH of the environment. LuCI is generated from sucralfate, an FDA-approved drug that selectively coats gastric ulcers. With the engineered modifications, LuCI forms a continuous coating on healthy mucosa. When exposed to gastrointestinal fluid, LuCI forms a sticky paste that binds to the mucosa and forms a barrier coating to block nutrient contact, mimicking the effects of RYGB. In our recently published work, we reported its single dose transient effects on lowering post prandial glucose responses in rodent models^22^, suggesting a therapeutic role in diabetes in a non-invasive manner (***Figure 1***).

**Figure 1.**
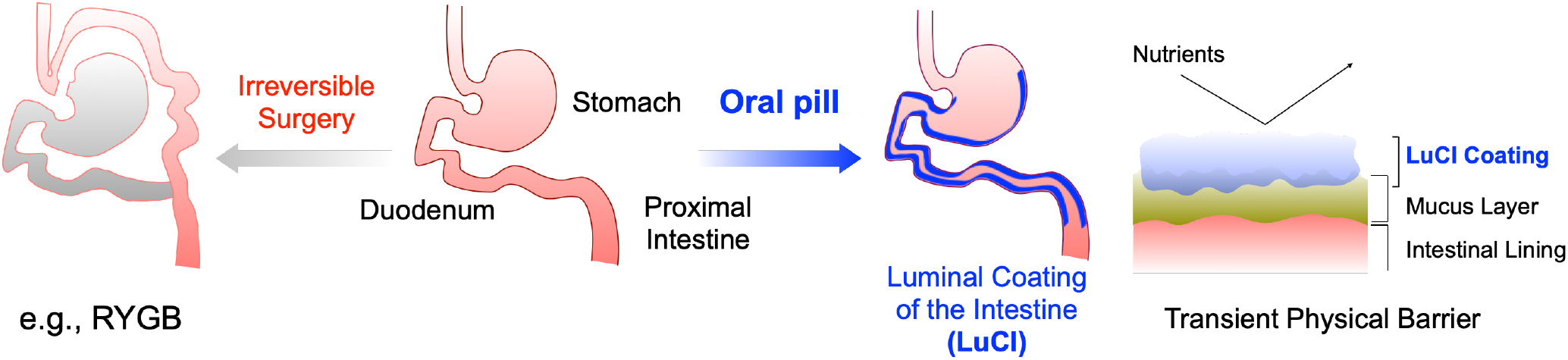
Representative cartoon demonstrating the oral administration luminal coating of intestine as an alternative to bariatric surgeries.

In this study, we designed a chronic daily dosing experiment to elicit LuCI’s effect on weight, hormones involved in glucose homeostasis as well as physiological energy balance in our high fat diet fed murine model. Namely, we hypothesized that LuCI improves glucose homeostasis, by acting as a transient intestine barrier coating, hence it could be utilized as a non-invasive therapeutic approach.

## Methods

### Experimental animals

Male Sprague-Dawley rats were purchased at 8 weeks of age from Charles River Laboratories (Wilmington, MA) and used in all experiments. Once arrived, rats were co-housed in the experimental room with controlled temperature and humidity. Animals were maintained on a 12h:12h dark:light cycle (light from 10pm to 10am and allowed *ad libitum* access to food and water unless otherwise specified. All experiments were performed in accordance with guidelines approved by the Institutional Animal Care and Use Committee at the Brigham and Women’s Hospital.

### Animal model

After 7 days of acclimatization on normal chow, animals were randomly assigned to vehicle or treatment group. Rats were then allowed *ad libitum* access to a high fat diet (HFD) (ResearchDiets, MA, 60% calories from fat, #D12492) throughout the experiment. Concordant to the start of HFD, treatment rats received a daily oral gavage of 450 g/kg rat LuCI 30 mins prior to the room’s dark cycle and start of their usual peak feeding time. Control group rats received an equivalent volume of normal saline.

### Formulation of LuCI

LuCI was fabricated using the method described in the previous study. Briefly, sucralfate was first treated with acid (0.6N HCl solutions) to form a viscous sticky paste that was further combined with ethanol and vortexed resulting in a white particle suspension. The suspension was then dried to remove the solvent and the dried particles were further ground to form a white powder. For a precise control on the dosing, LuCI was hydrated in a saline solution (0.9% w/v) before dosing and loaded in a syringe for gavage to rats.

### Body weight, food intake and stool collection

Body weight was monitored daily for 35 days at 10am using the same electronic digital scale. Food intake and stool output were measured once every 24 hours every 7^th^ day. On the 7^th^ day, a metal rack was placed at the bottom of the cage to prevent coprophagy. After 24 hours, food intake was determined by weighing the amount of food remaining from the previous day’s allowance. The metal rack was removed, and all visible stool pellets were collected and weighed. Stool pellets samples were stored at -80 °C for further analysis with bomb calorimetry.

### Functional glucose measurements

On the 21^st^ day of experiments, fasting glucose was measured. At 5 weeks, 2 sets of oral glucose tolerance tests (OGTT) and 1 insulin tolerance test (ITT) were carried out with at least 3 days between each test. For all the experiment, rats were fasted for 12 hours (10pm to 10am) prior to measurements. The 2 OGTTs were designed to demonstrate if the LuCI’s acute efficacy is still retained after 5 weeks of administration. Namely, in the first OGTT, the rats are deprived from any treatment for 24 hrs before OGTT as a baseline, and then in the second OGTT, the rats were gavaged with LuCI (for LuCI Group) or saline (for Control Group) 1hr prior to OGTT. In the first OGTT, animals were not given 24hrs with any LuCI (LuCI group) or normal saline (Control group). On the day of OGTT, animals were gavaged 2mg/g of oral D-Glucose (Sigma-Aldrich, St. Louis, MO). Baseline serum glucose levels were measured from a tail prick prior to glucose gavage and subsequently at 15, 30, 60, 90 and 120min with a OneTouch Glucometer (Life technologies, 252 San Diego, CA). Animals were then returned to their normal experimental routine for 2 days to recover. After 48 hours, we then sought to examine LuCIs acute on chronic effect, whereby a dose of LuCI or saline were administered 2 hours prior to OGTT.

To perform ITT, Actrapid HM human insulin (Eli Lily and Company, Indianapolis, IN)) at 0.6U/kg body weight was injected intraperitoneally (IP). Tail blood glucose was measured using OneTouch Glucometer (Life technologies, San Diego, CA) expressed in mg/dL. Baseline glucose was measured for each animal prior to insulin instillation with subsequent measurements at 15, 30, 60, and 90 minutes.

### Systemic blood collection by jugular vein catheterization and tissue harvest

Following 5 weeks of treatment, all rats were sacrificed, and tissues harvested at between 9am and 11am under general anesthesia using isoflurane (1-3% in oxygen). Animals were food deprived with *ad libitum* access to water for 12 hours prior to sacrifice unless specified otherwise. To ensure the data collected reflects the chronic treatment effect, the last dose of LuCI and normal saline was gavaged 24 hours prior to the day of harvest.

To allow the measurements of hormones following a glucose challenge, the duodenum was catheterized, and 2mg/kg D-Glucose (Sigma-Aldrich, St. Louis, MO) infused into the alimentary tract over 5 mins using a syringe driver (#NE 500, New Era Pump Systems, NY). A silastic catheter (0.02 in ID, Dow Corning, Midland, MI) was advanced within the left jugular vein into the right atrium, allowing sampling of mixed systemic venous blood at 0, 15, 30, 60 and 90 minutes time points. Blood samples were collected in previously prepared tubes containing chilled EDTA-containing Dipeptidyl Peptidase IV (DPPIV) inhibitor (10ul/ml blood; Millipore, Billerica, MA), Aprotinin (50ul/ml, Sigma-Aldrich, MA), and Pefabloc (10ul/ml, Sigma-Aldrich, MA). This was further centrifuged at 4000G for 10min at 4°C with plasma collected and stored at -80 °C for metabolic hormone assay. Liver and intestinal specimens were flash frozen in liquid nitrogen and stored at -80 °C until required.

### Biochemical analysis

The serum aliquots taken at the time of sacrifice were analyzed for insulin, GIP, GLP-1, ghrelin and leptin content using Luminex xMAP technology that is based on a multiplex ELISA-type fluorescence assay involving magnetic microspheres. Analysis was carried out on a Luminex 200 platform (Luminex, Austin, TX) using a Milliplex Rat Gut Hormone Panel 96-well plate assay as described by the manufacturer (#RMHMAG-84K-06; EMD Millipore Corporation, Billerica, MA). Data was collected with MAGPIX with xPONENT software (Luminex, Austin, TX).

### Bomb calorimetry

The energy content of feces collected at week 5 was determined by using an automatic Isoperibol 6725EA Semimicro Calorimeter and model 1109A Oxygen Combustion Bomb (Parr Instrument Co, Moline, IL). Detailed established methodology have been previously described^23^. In brief, the calorimeter was calibrated with benzoic acid to standardize the energy equivalent value. Fresh wet fecal pellet weight was measured to 0.0001g on an analytical balance as “wet weight”, then desiccated in an open microtube to dry in an oven at 600°C for 48 hours and reweighed as “dry weight”. The pellet was then placed in the oxygen bomb connected to a fuse wire and ignition unit surrounded by 2000ml distilled water. The heat produced by the combustion of the pellet was detected by the rise of water temperature. The energy content of the pellet was then calculated by comparing to the energy equivalent value. The final data obtained was expressed as calories per gram of dry pellet weight. Each sample was run in duplicate. Calorie ingested was determined by the amount of food ingested and the caloric value per gram of HFD (#D12492 at 5.21kcal/g). The value for caloric absorption and apparent digestibility of the food ingested were calculated as described^23^.

### Nuclear magnetic resonance spectroscopy (NMR) analysis

NMR spectra were acquired on a Vantera® Clinical Analyzer, a 400 MHz NMR instrument, from EDTA plasma samples as described for the NMR LipoProfile® test (Labcorp, Morrisville, NC). The NMR MetaboProfile analysis, using the LP4 lipoprotein profile deconvolution algorithm, reports lipoprotein particle concentrations and sizes, as well as concentrations of metabolites such as total branched chain amino acids, valine, leucine, and isoleucine, alanine, glucose, citrate, total ketone bodies, β-hydroxybutyrate, acetoacetate, acetone. The diameters of the various lipoprotein classes and subclasses are: total triglyceride-rich lipoprotein particles (TRL-P) (24-240 nm), very large TRL-P (90-240 nm), large TRL-P (50-89 nm), medium TRL-P (37-49 nm), small TRL-P (30-36 nm), very small TRL-P (24-29 nm), total low density lipoprotein particles (LDL-P) (19-23 nm), large LDL-P (21.5-23 nm), medium LDL-P (20.5-21.4 nm), small LDL-P (19-20.4 nm), total high density lipoprotein particles (HDL-P) (7.4-13.0 nm), large HDL-P (10.3-13.0 nm), medium HDL-P (8.7-9.5 nm), and small HDL-P (7.4-7.8 nm). Mean TRL, LDL and HDL particle sizes are weighted averages derived from the sum of the diameters of each of the subclasses multiplied by the relative mass percentage. Linear regression against serum lipids measured chemically in an apparently healthy study population (n=698) provided the conversion factors to generate NMR-derived concentrations of total cholesterol (TC), triglycerides (TG), TRL-TG, TRL-C, LDL-C and HDL-C. NMR-derived concentrations of these parameters are highly correlated (r ≥0.95) with those measured by standard chemistry methods. Details regarding the performance of the assays that quantify BCAA, alanine and ketone bodies have been reported. While these NMR assays have been analytically validated for use with human specimens, full analytical validation studies have not been performed in rodent specimens.

### Statistical analysis

Data are expressed as mean ± SEM. Statistical analysis was carried out using GraphPad Prism (version 8.4.2) and Student’s t-tests were used to compare two groups of continuous variables. Statistical tests used and P-values are mentioned in the figure legends. P value less than 0.05 was considered statistically significant.

## Results

### LuCI induces weight loss and lowers fasting glucose on HFD

The rats administered with daily LuCI showed a significant weight loss compared to the Control Group (***Figure 2A***). Control rats increased their body weight by 40.3% (257.8 ± 1.58g to 361.8 ± 5.82g) during the 5-week experimental period. In contrast, animals gavaged with LuCI gained only 32.1% (257.3 ± 2.27g to 339.8 ± 7.90g, P<0.05, ***Figure 2B***) over the 35 days and weighed 8.3% less than the control group. Measurements of food intake, however, did not reveal any differences between the 2 groups throughout the experiment (***Figure 2C***). The LuCI Group rats also showed lowered fasting glucose levels compared to the Control Group. Comparisons at 3 weeks showed that LuCI rats had a 21.7% reduction in fasting glucose levels compared to control group (77.6 ± 3.8mg/dl vs. 99.1 ± 2.7mg/dl, P<0.001, ***Figure 2D***). To determine whether chronically treated LuCI animals continued to respond to the oral coating agent, oral glucose tolerance testing (OGTT) was performed after 5 weeks and showed that LuCI lowered peak glucose level by 17.8% compared to normal saline group, (systemic glucose level at 60 min 128.3mg/dl vs. 156.1mg/dl, P<0.01, ***Figure 2E***) and significantly lowered area under the curve (AUC) (14225mg/dl vs. 16057mg/dl, P<0.001, ***Figure 2F***). We then repeated the experiments on another day, performing OGTT 24-hours after LuCI administration. At this time point, there was no changes were noted in OGTT parameters (Figure 2 E and F). Collectively, these data shows that the coating effect of LuCI in glucose handling appeared to be transient, and durable. Further more, once daily dosage of the drug had overall weight and metabolic benefits.

**Figure 2.**
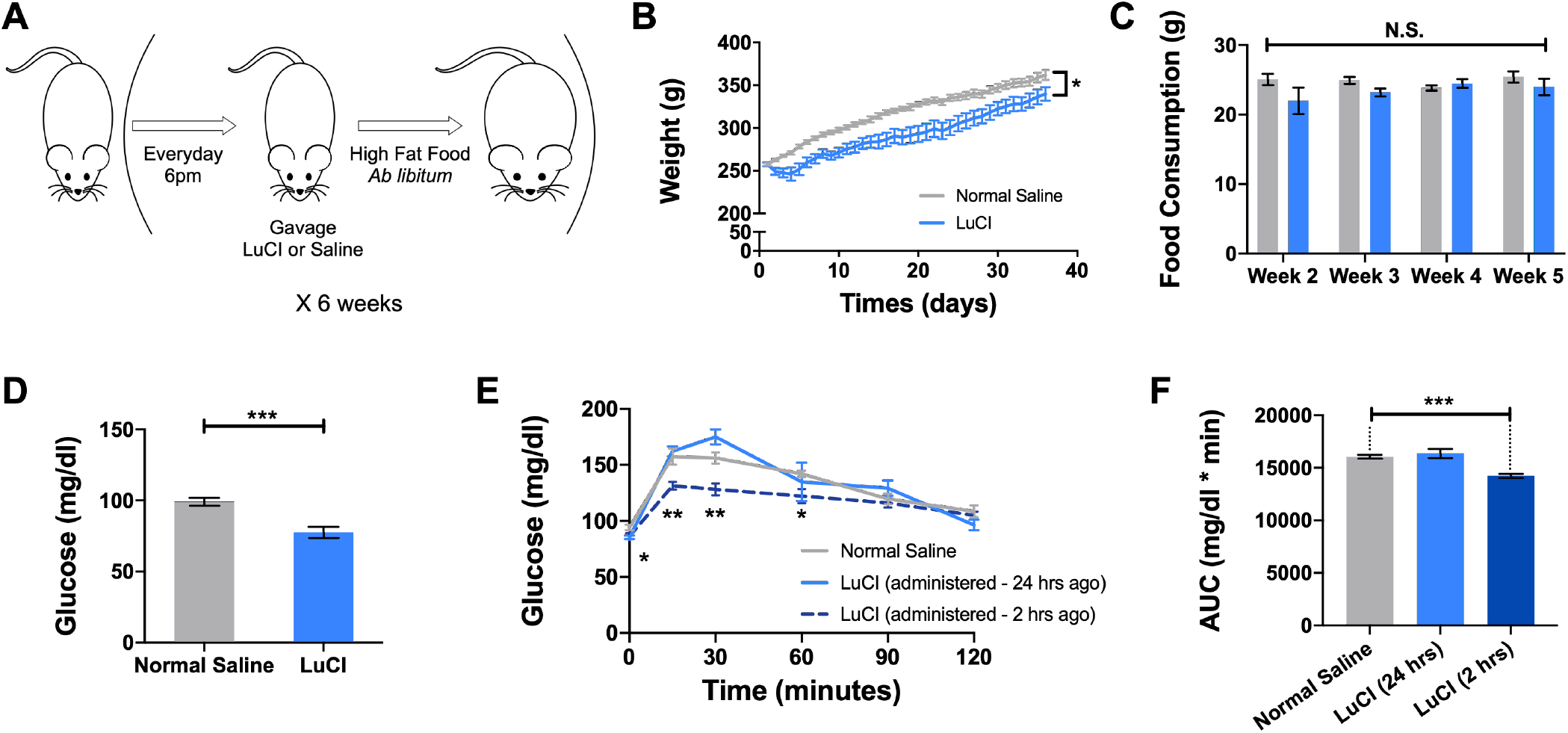
LuCI protects from weight gain and lowers fasting glucose on HFD. (A) Schematic representation of the experimental design. (B) Weight over 35 days of LuCI and vehicle oral gavage. (C) Food intake over 24 hours for each week. (D) Mean fasting glucose levels 3 weeks after beginning of treatment. (E) Oral glucose tolerance testing at 5 weeks. (F) Area under the curve (AUC). (B-F) n = 8, unpaired t test. *P<0.05, **P<0.01, ***P<0.001. Data are presented as mean values ± SEM.

### LuCI mitigates insulin resistance and lowers HOMA-IR

We next measured the effects of LuCI on insulin tolerance after 5 weeks of treatment. An insulin tolerance test at 5 weeks revealed that control rats had an impaired insulin tolerance compared to the LuCI-treated rats (***Figure 3A***). Although there was no difference in insulin secretion 15 – 120 min following a glucose challenge (***Figure 3B and 3C***), fasting insulin was 5.3-fold lower in the LuCI group as compared to the normal saline group (1133.0 ± 48.8pg/ml vs. 212.2 ± 29.4pg/ml, P<0.001) Likewise, the LuCI group demonstrated a significant 6.1-fold reduction in HOMA-IR as compared to the control group (6.66 ± 1.08 vs. 0.08 ± 0.18, P<0.001, ***Figure 3D***).

**Figure 3.**
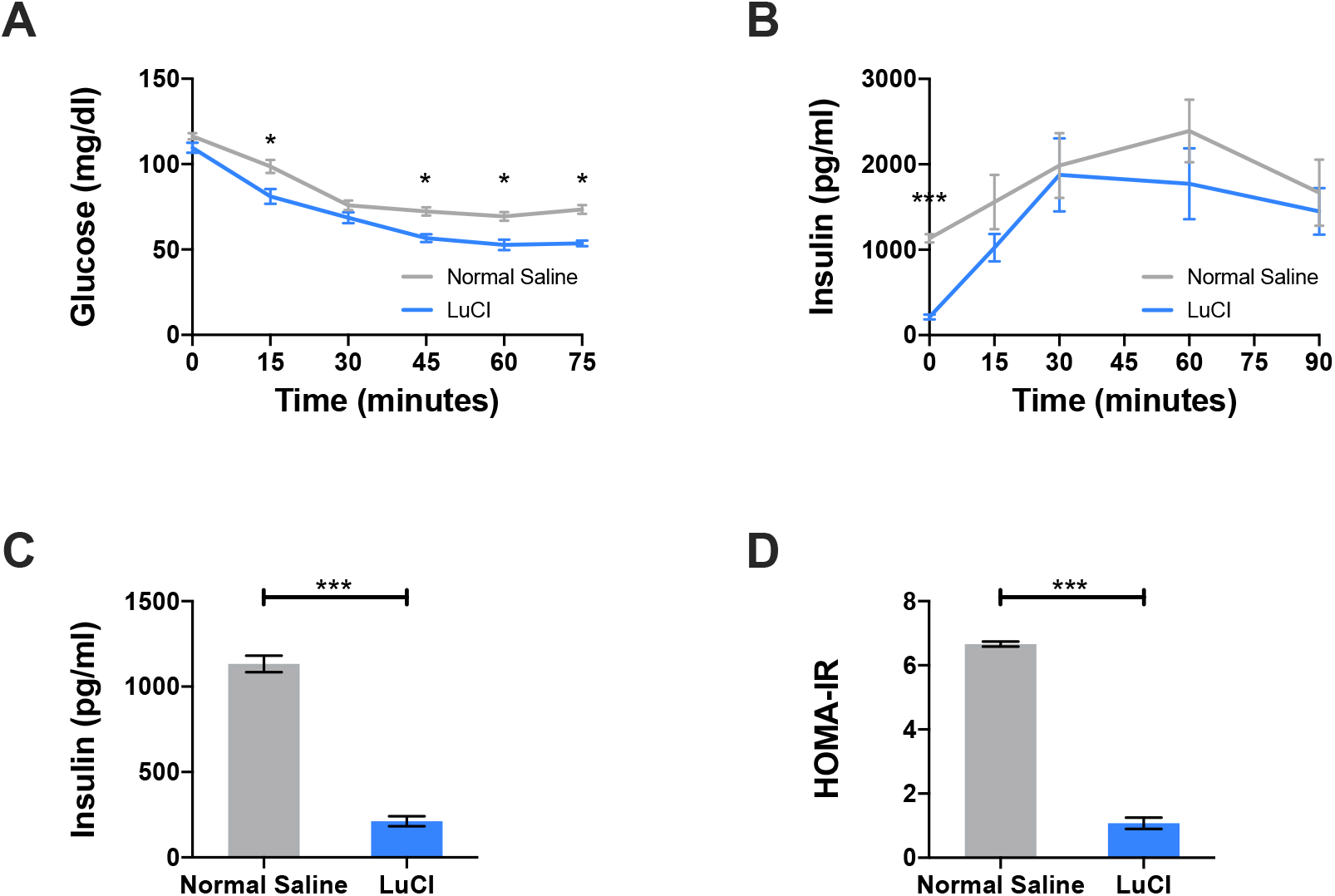
LuCI treatment mitigates insulin resistance and lowers HOMA-IR on HFD. (A) Tail blood glucose values during an insulin tolerance test (ITT) for LuCI-treated and normal saline rats at 5 weeks. (B-D) Systemic circulating insulin levels for LuCI and NS rats in response to a duodenal glucose infusion. (B) Insulin levels at time points indicated. (C) Systemic fasting insulin level. (D) HOMA-IR. (A-D) n = 7-8, unpaired t test. *P<0.05, ***P<0.001. Data are presented as mean values ± SEM.

### LuCI reduces circulating incretins during fasting and improves GLP-1 and GIP responses following a glucose challenge

To examine whether LuCI treatment over 5 weeks exert an effect on hormones involved with glycemic control, we analyzed the circulating plasma hormone levels following a duodenal glucose infusion over specific time points. In accordance with the acknowledged post RYGB physiology, LuCI rats had a significantly increased response in circulating incretins such as GLP-1 and GIP starting at 15 min after the glucose challenge. Their perspective AUC were also higher in LuCI group (***Figure 4A and 4B*)**.

**Figure 4.**
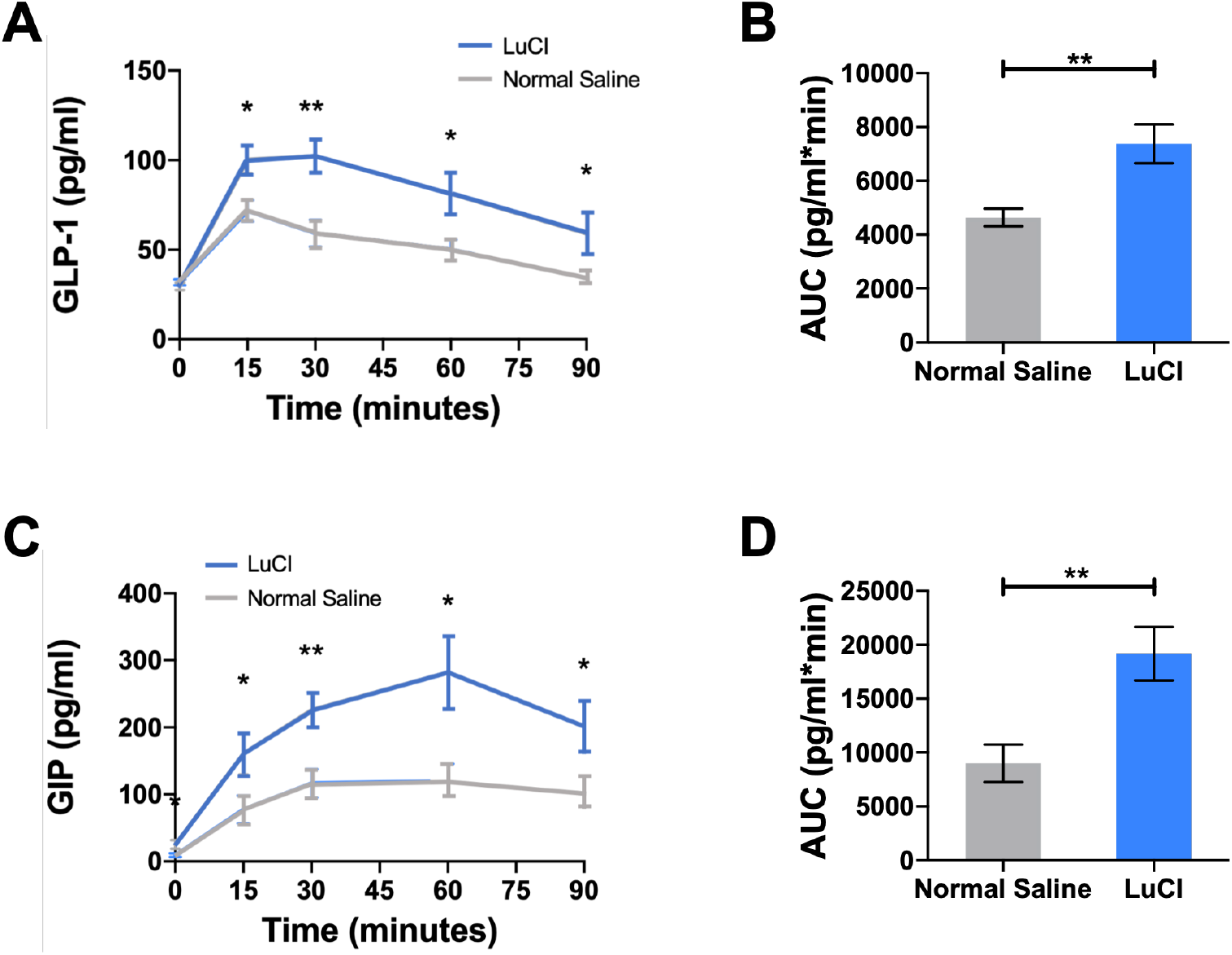
LuCI reduced circulating incretins during fasting and improved GLP-1 and GIP response following a glucose challenge. (A-D) Circulating plasma hormone levels measured 5 weeks following daily gavage of LuCI or normal saline. (A) GLP-1 levels at time points indicated. (B) Area under the curve (AUC) for GLP-1 in (A). (C) GIP levels at time points indicated. (D) Area under the curve (AUC) for GIP-1. (A-D) n = 7-8, unpaired t test. *P<0.05, **P<0.01. Data are presented as mean values ± SEM.

### LuCI alters appetite regulating hormones

The fasting ghrelin and leptin levels in the systemic circuit were shown to be 2.0-fold (57.3 ± 8.31pg/ml vs 29.1 ± 4.29pg/ml, P<0.01, ***Figure 5A***) and 1.8-fold (553.3 ± 95.58pg/ml vs 303.0 ± 52.25pg/ml, P<0.05, ***Figure 5B***) respectively lower after 5 weeks of LuCI as compared to the control group. In the following calorimetric analysis, we measured the amount of calories consumed by the amount of food ingested and the caloric value per gram of diet (LuCI: 62.5.1 ± 3.08kcal vs NS: 64.2 ± 2.74kcal, P=0.71, ***Figure 6A***).

**Figure 5.**
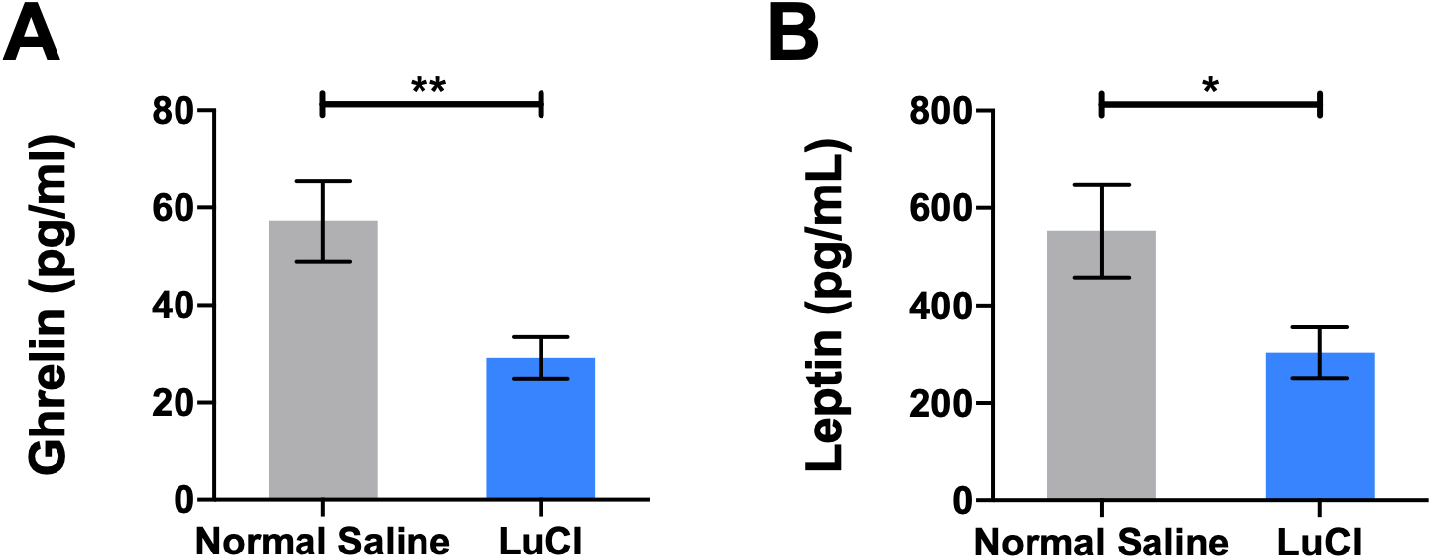
LuCI alters fasting levels of appetite-regulating hormones. (C) Fasting ghrelin levels. (D) Fasting leptin levels. (A-B) n = 7-8, unpaired t test. *P<0.05. Data are presented as mean values ± SEM.

**Figure 6.**
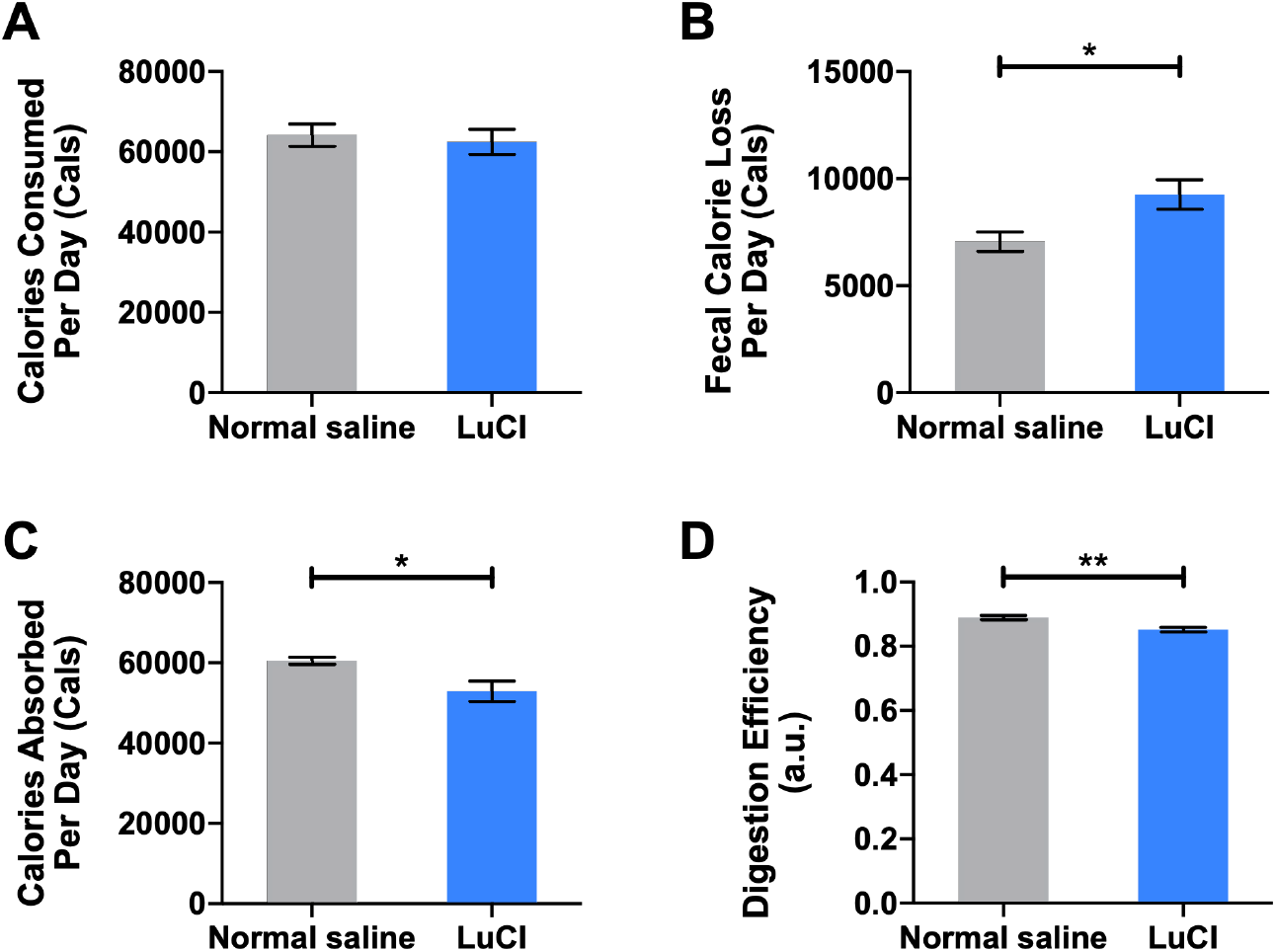
LuCI reduces calorie absorption in the gut with no difference in calorie consumption. (A-D) Fecal bomb calorimetry analysis 5 weeks following daily gavage of LuCI or normal saline. (A) Calories consumed per day. (B) Fecal caloric loss per day. (C) Calories absorbed per day. (D) Digestion efficiency. (A-D) n = 4, unpaired t test. *P<0.05, **P<0.01. Data are presented as mean values ± SEM.

Using bomb calorimetry, we further examined the fecal energy content and calculated the differences in energy absorption between the groups. We found that despite the desiccated fecal pellets from LuCI rats tend to be heavier (LuCI: 21.3mg vs NS: 11.4g, P<0.05, ***Supplementary figure 1B***), their caloric density was found to significantly less (LuCI: 4.1kcal vs NS: 4.7kcal, P<0.01, ***Supplementary figure 1C***). However, although comprised of a lower caloric density, the combined effects of increased fecal production resulted in a significantly increased total fecal energy output 5 weeks after LuCI treated rats compared with control cohort (LuCI: 9.13 ± 0.69kcal vs NS: 7.1 ± 0.45kcal) (***Figure 6B***). Together with the number of calories consumed considered, we can deduce that LuCI treated rats have a 14.3% lower caloric absorption per day (LuCI: 52.0 ± 2.42kcal vs NS: 60.6 ± 0.91kcal) (***Figure 6C***) and are therefore less efficient in nutrition digestion (***Figure 6D***).

### LuCI treatment lowered the total serum cholesterol level, especially contributed by the lowered serum LDL levels

Using the NMR-based serum lipid panel analyses, LuCI treated rats showed lowered serum cholesterol levels (66.0 mg/dL in LuCI Group vs 88.5 mg/dL in Control Group, p=0.0214 in Student t-test, ***Figure 7A***), and especially lowered circulating LDL concentration compared to the Control Group rats (47.75 mg/dL in LuCI Group vs 73.0 mg/dL in Control Group, p=0.0082 in Student t-test, ***Figure 7B***). More detailed analyses showed that the subclass of large LDL particles was specifically lowered 2.6-fold compared to the Control Group (233.8 mmol/L in LuCI Group vs 616.3 mmol/L in Control Group, p=0.0004 in Student t-test, ***Figure 7C***) and also the lower average size of the LDL particles (21.59 nm in LuCI Group vs 22.13 nm in Control Group, p=0.0010 in Student t-test, ***Figure 7D***), suggesting the potential benefit on cardiovascular risks (also see ***Supplementary Figure 2-7*** for the full lipid panel).

**Figure 7.**
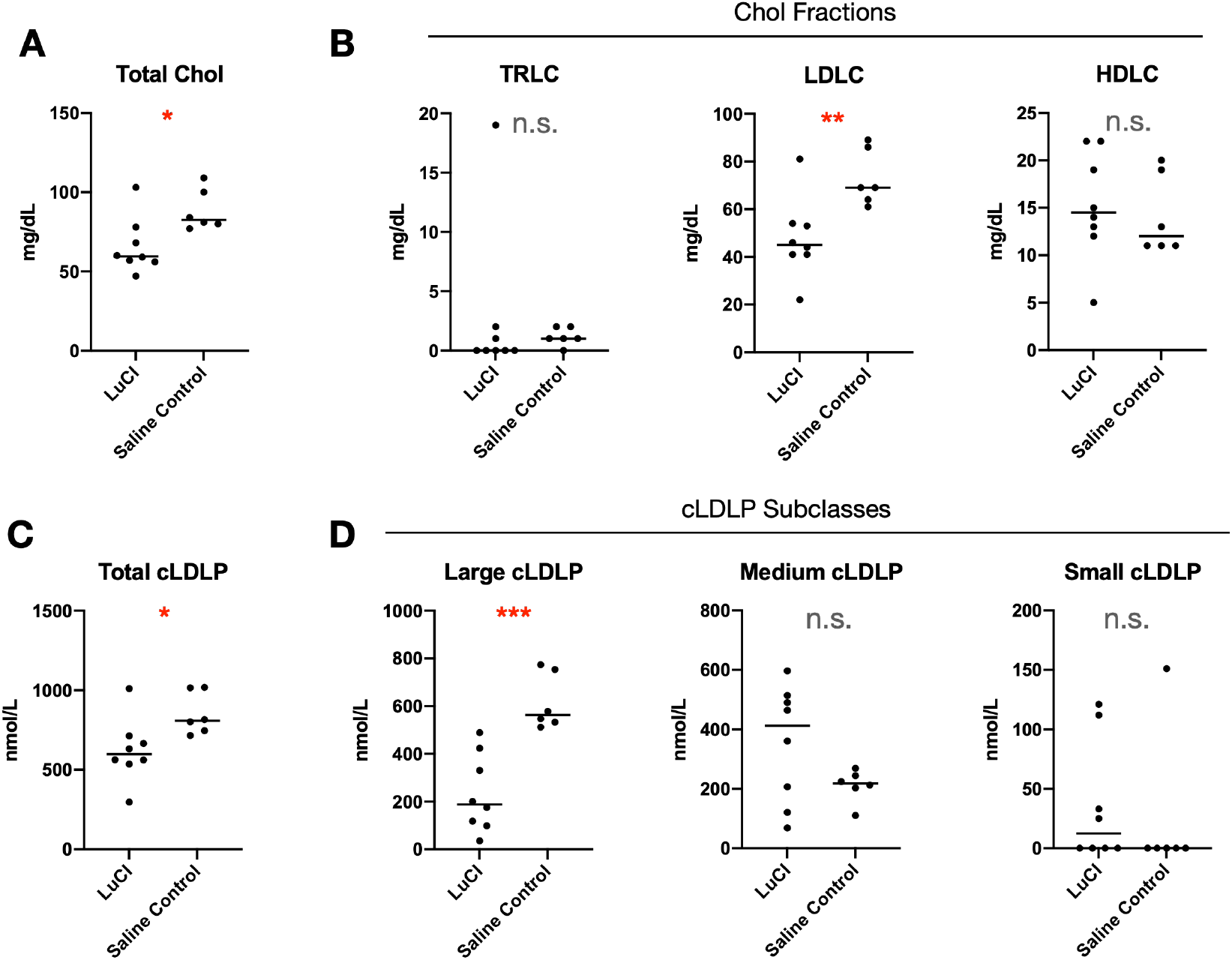
Serum lipid panel analysis for LuCI Group and Control Group after 5 weeks. (a) Total cholesterol concentration. (b) Analysis of cholesterol fractions (TRLC: triglyceride-rich lipoprotein concentration, LDLC: low density lipoprotein concentration, HDLC: high density lipoprotein concentration). (c) Total circulating LDL particle concentration. (d) Subclass analysis of (c).

## Discussion

Foregut exclusion plays a pivotal role in the metabolic benefits that occur following RYGB. It has been demonstrated that through bypassing proximal intestines, glucose tolerance can be improved independently of weight loss and food intake^20,21,24,25^. The underlying mechanisms and their relevance to the improved metabolic profile have been extensively investigated over the past decade. Proposed pathways include normalization of metabolic hormone secretions such as peptide YY (PYY)^26,27^, gastric inhibitory peptide (GIP)^28^ and glucagon-like peptide-1 (GLP-1)^29,30^, stimulation of glucose sensing transporters^15^ and vagal afferent nerves^31–33^, as well as changes in gut microbiome^29,34,35^ and bile acid levels^13,34^. The combination of these physiological alterations reduces hunger sensations and enhances tissues’ energy expenditure and subsequently improves insulin sensitivity and glucose tolerance.

Current research exploring potential mechanisms of anti-diabetic effect of RYGB surgery has hilight several potential mechanisms, with exclusion of the proximal bowel from nutrient exposure being an important step. This has been the drier behind our work to create a transient coating ot eh bowel to replicates the metabolic effect of surgery in a less invasive fashion. Our design criteria included an oral treatment that would create a transient barrier only to avoid long term complication and easy reversibility. Our work has led to creation of LuCI which has been in shown in our previous publications to significantly lower the OGTT response (47% reduction in iAUC), through a local gut barrier function and without any systemic effect.

We have also shown that a single dose of LuCI, an orally administered therapy that modulated the nutrient contact with foregut mucosa, lowered glucose responses indicating a potential therapeutic utility in T2D^22^. In the present study, we aim to demonstrate the long-term effects of LuCI on weight and insulin resistance. Studies have shown that although almost half of the population in the United States fit the criteria for use of anti-obesity pharmacotherapy, only 2% of those receive such treatment^39^. Principally, it is therefore crucial to determine LuCI’s role in the early treatment of obesity and prevention of its associated metabolic diseases progression. To assess its preventative properties, we established a diet-induced obesity rodent model in conjunction with a daily, weight-dependent dose of LuCI over the course of 5 weeks. We inferred its long-term therapeutic outcomes that has reproduced similar changes observed in human RYGB patients^12,14,24,29^, including a significant difference in weight, improvement in insulin sensitivity, and increases in plasma GLP-1, GIP hormones. Furthermore, potential for improvement in cardiovascular health was highlighted by the lowering of cholesterol and LDL levels. The changes seen are similar to that seen in RYGB.

When exploring potential mechanism for these beneficial changes, we saw a lowering in plasma ghrelin levels which has also been reported after RYGB. Although this did not translate to a lowering of daily food intake, it is possible that the decrease was small and our current study underpowered to detect it, and larger rodent groups with more frequent daily food intake is required to assess this. It is also important to note that no clinical diarrhea was observed throughout the experimental period, however the weight of stool output was higher in the *LuCI* group and this lead to an increase in stool calorie output which likely contributed to the weight loss and improvement in glucose homeostasis. Similar results have been seen in rodent models of RYGB.

Notably, the concept of foregut exclusion which mimics RYGB’s ability to ameliorate weight gain and T2D is not completely novel. Over the years, scientists have been eager to replicate the effects of RYGB with innovative techniques and devices that possess minimal surgical risks, and with effects that can be potentially reversible. The development of duodenal-jejunal bypass sleeve (DJBS) demonstrated a functional device that prevented contact between chime and the proximal intestinal mucosa. The endoscopically delivered device showed promising results in remitting T2D in patients and a 12% weight loss compared with sham controls or dietary intervention alone^40–43^. This further validates the concept that isolation of the proximal intestines from nutrient exposure can lead to dramatic improvements in weight and T2D. However, DJBS requires endoscopic annual device removal hence harbours technical risks and its first pivotal US ENDO trial was initially halted in 2015 due to increased risk of hepatic abscess formation^44^. Further trials and studies are therefore underway to determine its safety and efficacy.

As an oral administered agent, LuCI has been shown to be safe, effective, and non-invasive. It has been bio-engineered and altered from the FDA-approved drug, sucralfate, to bound with water and form an effective mucosal coating independent of pH^22^. This coating effect, however, had been shown to be non-systemic and transient while providing a temporary proximal bowel exclusion effect^22^. In its powder form, LuCI can be manufactured within a capsule and delivered to a destinated part of gastrointestinal tract. These unique properties may allow early intervention in obesity and prevent progression of T2D while allowing patients to take control of their treatment according to their individual lifestyle.

Our study has several limitations. The use of diet induced obesity rodent models is unable to completely replicate human obesity and its associated T2D disease process, however, is essential in target validation and the early pre-clinical proof of concept stage. Furthermore, although we observed both safety and efficacy with LuCI at a dose of 450mg/kg daily in rats, further experiments will be required to elicit the optimal dosage for humans. Future animal studies on whether LuCI alters other parameters of mechanisms previously mentioned, such as gut microbiome and bile acids secretion will also allow us to further understand the modality in which LuCI function. Eventually, this will allow us to utilize the translational value of animal experiments to human studies so to confirm our findings and hypothesis.

In summary, we have demonstrated that LuCI, an orally administered intestine barrier coating, can ameliorate weight gain, and improve insulin sensitivity in a long-term DIO rat model. At 5 weeks, LuCI animals weighted 8.3% less and had lower fasting glucose levels than Controls (77.6 ± 3.8 mg/dl vs. 99.1 ± 2.7 mg/dl, P<0.001). LuCI-treated animals had lower insulin and HOMA-IR. Post-prandially, LuCI group had increased GLP-1 and GIP secretion following a glucose challenge. Serum lipid analysis revealed lowered LDL and improved lipoprotein particles. There was no difference in food intake between the groups, but bomb calorimetry showed an increase in stool calories. Since LuCI’s effect is transient, and there is no systemic absorption of the drug with minimal safety concerns, LuCI has the potential to be beneficial for those who are overweight or obese. Future studies should consider LuCI as a new potential therapeutic strategy to limit weight gain and prevent the development of T2D.

## Supplementary Figures

**Supplementary figure 1.**
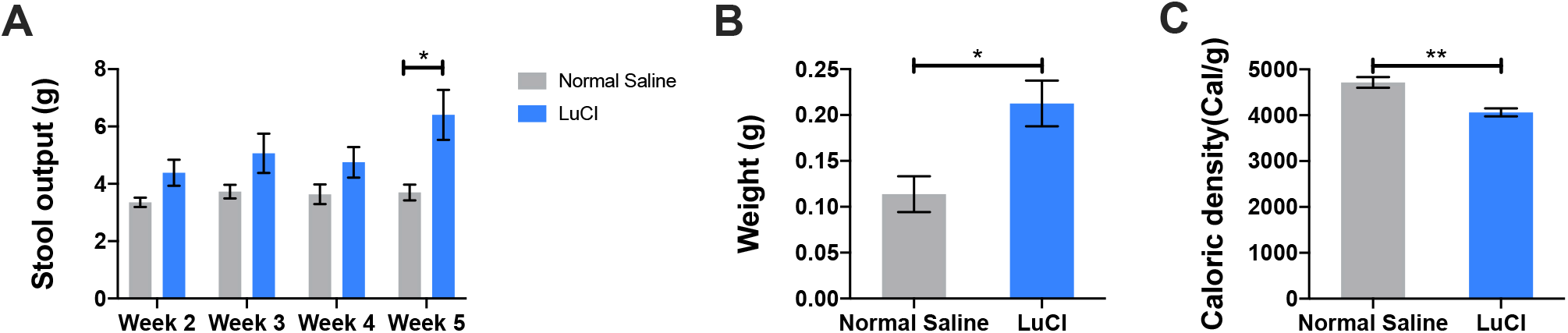
(A) Stool output in 24 hours. (B) Dry weight of fecal pellet (C) Calories density elicited from bomb calorimetry per gram of desiccated pellet. (A-C) n = 4, unpaired t test. *P<0.05, **P<0.01. Data are presented as mean values ± SEM.

**Supplementary Figure 2.**
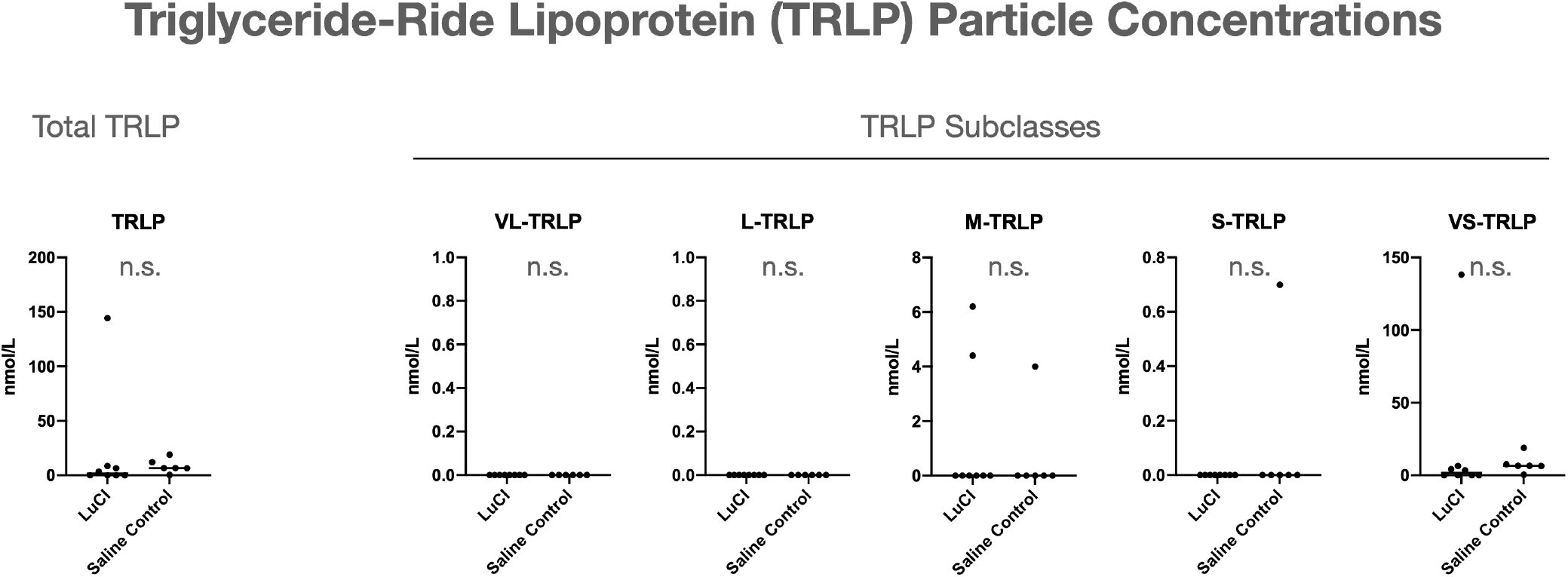
Serum triglyceride-ride lipoprotein (TRLP) particle concentrations of the rats treated with LuCI or saline for 6 weeks measured by NMR analysis (see ***Methods*** section for more details). VL-TRLP: Very Large TRLP, L-TRLP: Large TRLP, M-TRLP: Mid-size TRLP, S-TRLP: Small TRLP, VS-TRLP: Very Small TRLP.

**Supplementary Figure 3.**
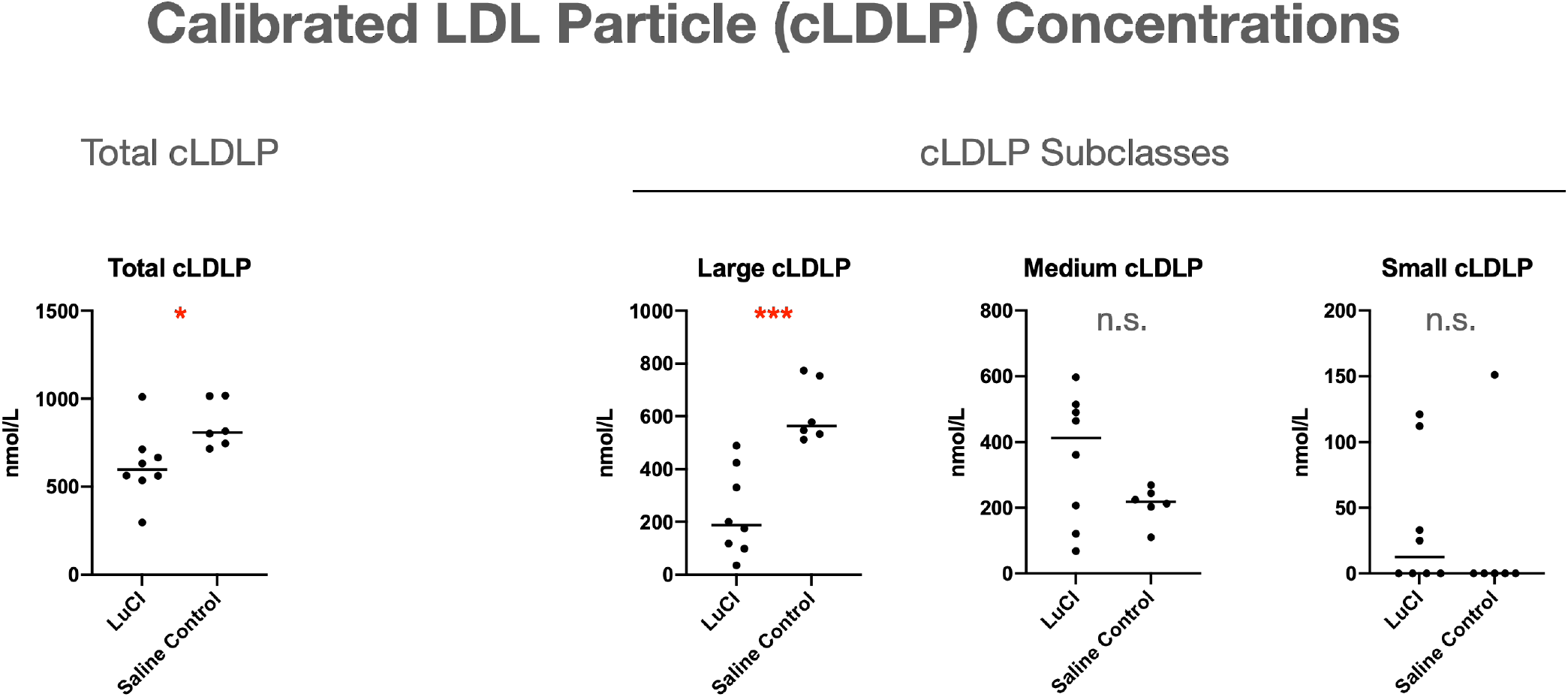
Serum calibrated LDL Particle (cLDLP) concentrations of the rats treated with LuCI, or saline for 6 weeks measured by NMR analysis (see ***Methods*** section for more details).

**Supplementary Figure 4.**
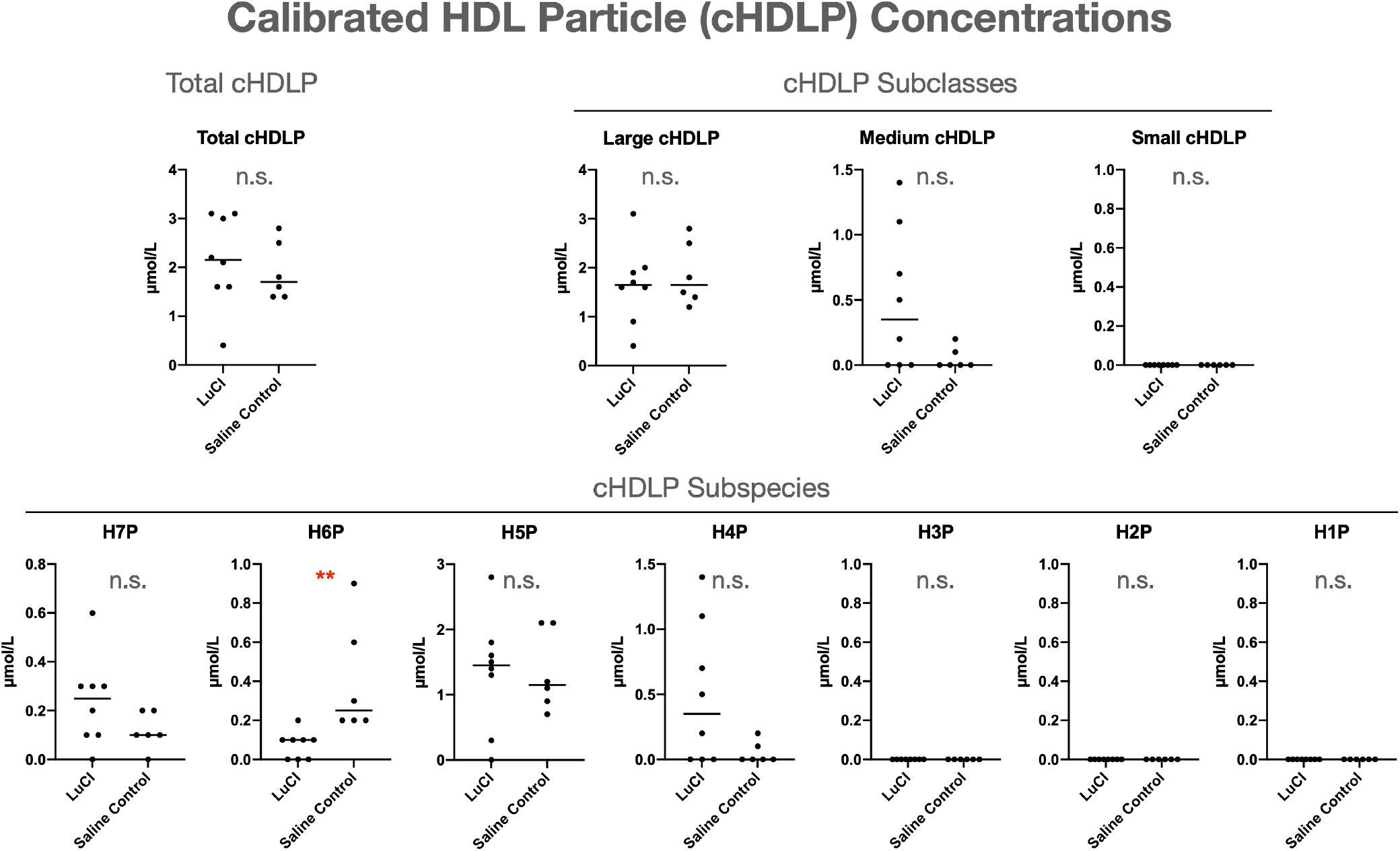
Serum calibrated HCL particle (cHDLP) concentrations of the rats treated with LuCI, or saline for 6 weeks measured by NMR analysis (see ***Methods*** section for more details).

**Supplementary Figure 5.**
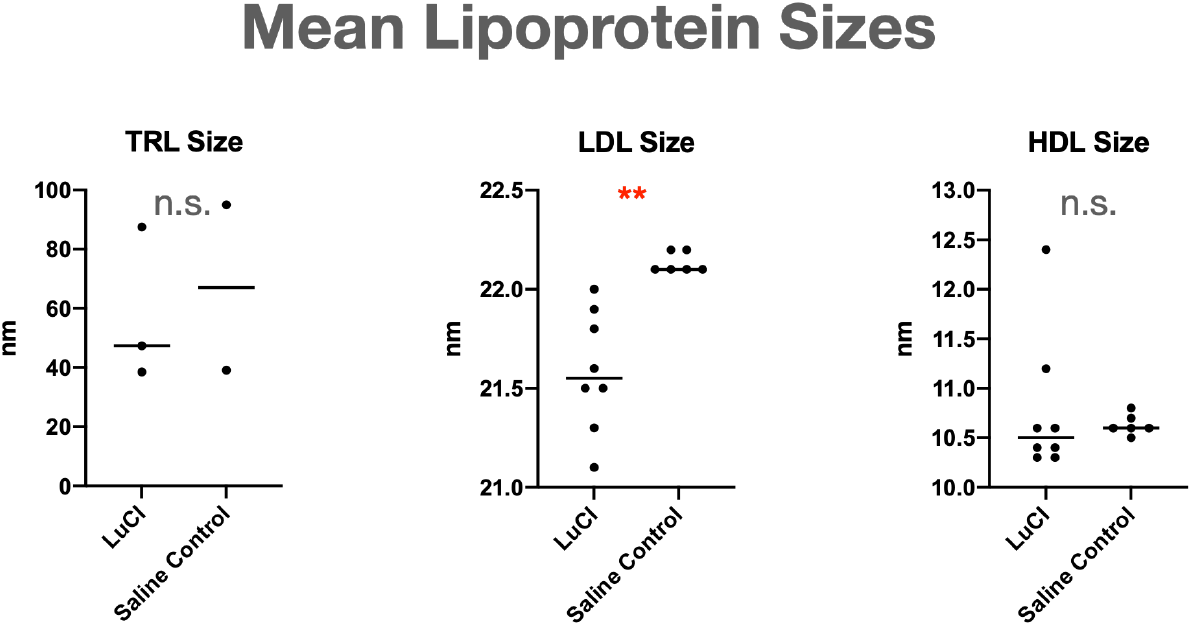
Serum mean lipoprotein sizes of the rats treated with LuCI or saline for 6 weeks measured by NMR analysis (see ***Methods*** section for more details).

**Supplementary Figure 6.**
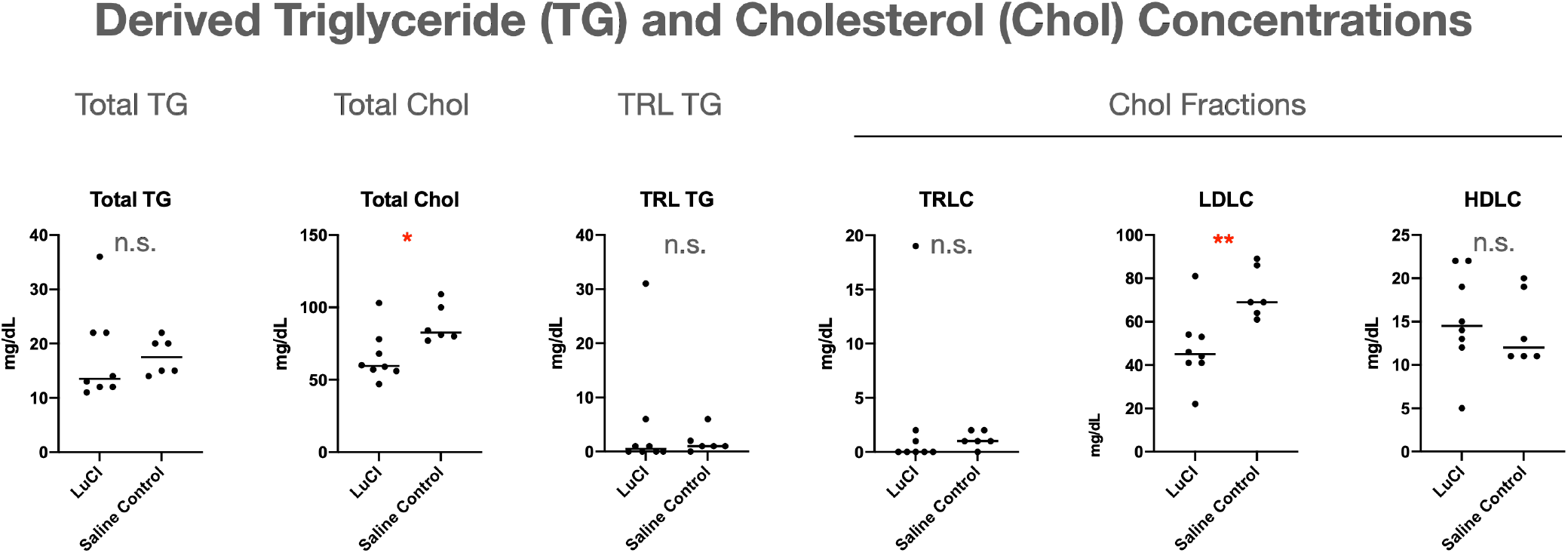
Serum derived triglyceride and cholesterol concentrations of the rats treated with LuCI or saline for 6 weeks measured by NMR analysis (see ***Methods*** section for more details).

## Author Contributions

- T.L., Y.L, J.M.K, C.S.M. and A.T. designed this work.
- J.M.K, and A.T. co-supervised this work.
- T.L., C.-Y.T., J.H. and Y.L. performed the animal in vivo experiments.
- T.L. processed the samples and measured the biomarkers.
- T.L. performed the statistical analyses.
- All co-authors read and approved this work.

## Acknowledgements

Authors thank Nancy Briefs in AltrixBio inc. for helpful comments and discussions. Authors also thank Margery Connelly in LabCorp, inc. for the lipidomics analysis.

## Reference

1. Colditz GA, Willett WC, Rotnitzky A, Manson JE. Weight gain as a risk factor for clinical diabetes mellitus in women. Ann Intern Med. 1995;122(7):481-486. doi:10.7326/0003-4819-122-7-199504010-00001

2. Willett WC, Dietz W, Colditz GA. Guidelines for healthy weight. N Engl J Med. 1999;341(6):427–434. www.nejm.org. Accessed April 21, 2020.

3. WHO. Fact Sheet: Obesity and overweight. https://www.who.int/en/news-room/fact-sheets/detail/obesity-and-overweight. Published 2020.

4. Schauer PR, Bhatt DL, Kirwan JP, et al. Bariatric surgery versus intensive medical therapy for diabetes - 5-year outcomes. N Engl J Med. 2017;376(7):641–651. doi:10.1056/NEJMoa1600869

5. Mingrone G, Panunzi S, De Gaetano A, et al. Bariatric-metabolic surgery versus conventional medical treatment in obese patients with type 2 diabetes: 5 Year follow-up of an open-label, single-centre, randomised controlled trial. Lancet. 2015;386(9997):964–973. doi:10.1016/S0140-6736(15)00075-6

6. Simonson DC, Halperin F, Foster K, Vernon A, Goldfine AB. Clinical and patient-centered outcomes in obese patients with type 2 diabetes 3 years after randomization to Roux-en-Y gastric bypass surgery versus intensive lifestyle Management: The SLIMM-T2D study. Diabetes Care. 2018;41(4):670–679. doi:10.2337/dc17-0487

7. Courcoulas AP, Belle SH, Neiberg RH, et al. Three-year outcomes of bariatric surgery vs lifestyle intervention for type 2 diabetes mellitus treatment a randomized clinical trial. JAMA Surg. 2015;150(10):931–940. doi:10.1001/jamasurg.2015.1534

8. American Society for Metabolic and Bariatric Surgery. Estimate of Bariatric Surgery Numbers - American Society for Metabolic and Bariatric Surgery. www.asmbs.org. 2014. https://asmbs.org/resources/estimate-of-bariatric-surgery-numbers https://asmbs.org/resources/estimate-of-bariatric-surgery-numbers http://asmbs.org/resources/estimate-of-bariatric-surgery-numbers.

9. Knop FK, Taylor R. Mechanism of metabolic advantages after bariatric surgery: It’s all gastrointestinal factors versus it’s all food restriction. Diabetes Care. 2013;36(SUPPL.2):287–291. doi:10.2337/dcS13-2032

10. Cummings DE. Endocrine mechanisms mediating remission of diabetes after gastric bypass surgery. Int J Obes. 2009;33:33–40. doi:10.1038/ijo.2009.15

11. Heshmati K, Harris DA, Aliakbarian H, Tavakkoli A, Sheu EG. Comparison of early type 2 diabetes improvement after gastric bypass and sleeve gastrectomy: medication cessation at discharge predicts 1-year outcomes. Surg Obes Relat Dis. 2019;15(12):2025–2032. doi:10.1016/j.soard.2019.04.004

12. Le Roux CW, Aylwin SJB, Batterham RL, et al. Gut hormone profiles following bariatric surgery favor an anorectic state, facilitate weight loss, and improve metabolic parameters. Ann Surg. 2006;243(1):108–114. doi:10.1097/01.sla.0000183349.16877.84

13. Bhutta HY, Rajpal N, White W, et al. Effect of Roux-en-Y gastric bypass surgery on bile acid metabolism in normal and obese diabetic rats. PLoS One. 2015;10(3):1–17. doi:10.1371/journal.pone.0122273

14. Bhutta HY, Deelman TE, le Roux CW, Ashley SW, Rhoads DB, Tavakkoli A. Intestinal sweet-sensing pathways and metabolic changes after Roux-en-Y gastric bypass surgery. Am J Physiol - Gastrointest Liver Physiol. 2014;307(5). doi:10.1152/ajpgi.00405.2013

15. Pal A, Rhoads DB, Tavakkoli A. Effect of portal glucose sensing on systemic glucose levels in SD and ZDF Rats. PLoS One. 2016;11(11):1–14. doi:10.1371/journal.pone.0165592

16. Liou AP, Paziuk M, Luevano JM, Machineni S, Turnbaugh PJ, Kaplan LM. Conserved shifts in the gut microbiota due to gastric bypass reduce host weight and adiposity. Sci Transl Med. 2013;5(178):178–219. doi:10.1126/scitranslmed.3005687

17. Sala P, Belarmino G, Machado NM, et al. The SURMetaGIT study: Design and rationale for a prospective pan-omics examination of the gastrointestinal response to Roux-en-Y gastric bypass surgery. J Int Med Res. 2016;44(6):1359–1375. doi:10.1177/0300060516667862

18. Thaler JP, Cummings DE. Minireview: Hormonal and metabolic mechanisms of diabetes remission after gastrointestinal surgery. Endocrinology. 2009;150(6):2518–2525. doi:10.1210/en.2009-0367

19. Madsbad S, Dirksen C, Holst JJ. Mechanisms of changes in glucose metabolism and bodyweight after bariatric surgery. Lancet Diabetes Endocrinol. 2014;2(2):152–164. doi:10.1016/S2213-8587(13)70218-3

20. Cummings DE, Overduin J, Foster-Schubert KE, Carlson MJ. Role of the bypassed proximal intestine in the anti-diabetic effects of bariatric surgery. Surg Obes Relat Dis. 2007;3(2):109–115. doi:10.1016/j.soard.2007.02.003

21. Rubino F, Forgione A, Cummings DE, et al. The mechanism of diabetes control after gastrointestinal bypass surgery reveals a role of the proximal small intestine in the pathophysiology of type 2 diabetes. Ann Surg. 2006;244(5):741–749. doi:10.1097/01.sla.0000224726.61448.1b

22. Lee Y, Deelman TE, Chen K, Lin DSY, Tavakkoli A, Karp JM. Therapeutic luminal coating of the intestine. Nat Mater. 2018;17(9):834–842. doi:10.1038/s41563-018-0106-5

23. Grobe JL. Comprehensive assessments of energy balance in mice. Methods Mol Biol. 2017;1614:123–146. doi:10.1007/978-1-4939-7030-8_10

24. Peterli R, Wölnerhanssen B, Peters T, et al. Improvement in glucose metabolism after bariatric surgery: Comparison of laparoscopic roux-en-Y gastric bypass and laparoscopic sleeve gastrectomy: A prospective randomized trial. Ann Surg. 2009;250(2):234–241. doi:10.1097/SLA.0b013e3181ae32e3

25. Hansen EN, Tamboli RA, Isbell JM, et al. Role of the foregut in the early improvement in glucose tolerance and insulin sensitivity following Roux-en-Y gastric bypass surgery. Am J Physiol - Gastrointest Liver Physiol. 2011;300(5). doi:10.1152/ajpgi.00019.2011

26. Patrício BG, Morais T, Guimarães M, et al. Gut hormone release after gastric bypass depends on the length of the biliopancreatic limb. Int J Obes. 2018:1–10. doi:10.1038/s41366-018-0117-y

27. Magouliotis D, Tasiopoulou V, Sioka E, Chatedaki C, Zacharoulis D. Impact of Bariatric Syrgery on Metabolic and Gut Microbiota Profile: a Systemic Review and Meta-analysis. Obes Surg. 2017;27:1345– 1357. doi:10.1007/s11695-017-2595-8

28. Palha AM, Pereira SS, Costa MM, et al. Differential GIP/GLP-1 intestinal cell distribution in diabetics’ yields distinctive rearrangements depending on Roux-en-Y biliopancreatic limb length. J Cell Biochem. 2018;119(9):7506–7514. doi:10.1002/jcb.27062

29. Mishra AK, Dubey V, Ghosh AR. Obesity: An overview of possible role(s) of gut hormones, lipid sensing and gut microbiota. Metabolism. 2016;65(1):48–65. doi:10.1016/j.metabol.2015.10.008

30. Hutch CR, Sandoval D. The role of GLP-1 in the metabolic success of bariatric surgery. Endocrinology. 2017;158(12):4139–4151. doi:10.1210/en.2017-00564

31. Blasi C. The Role of the Vagal Nucleus Tractus Solitarius in the Therapeutic Effects of Obesity Surgery and Other Interventional Therapies on Type 2 Diabetes. Obes Surg. 2016;26(12):3045–3057. doi:10.1007/s11695-016-2419-2

32. Stearns AT, Ashley SW, Balakrishnan A, Rhoads DB, Radmanesh A, Tavakkolizadeh A. Relative contributions of afferent vagal fibers to resistance to diet-induced obesity. Dig Dis Sci. 2012;57(5):1281–1290. doi:10.1007/s10620-011-1968-4

33. Malbert CH, Picq C, Divoux JL, Henry C, Horowitz M. Obesity-Associated alterations in glucose metabolism are reversed by chronic bilateral stimulation of the abdominal vagus nerve. Diabetes. 2017;66(4):848–857. doi:10.2337/db16-0847

34. Fouladi F, Mitchell JE, Wonderlich JA, Steffen KJ. The Contributing Role of Bile Acids to Metabolic Improvements After Obesity and Metabolic Surgery. Obes Surg. 2016;26(10):2492–2502. doi:10.1007/s11695-016-2272-3

35. Murphy R, Tsai P, Jüllig M, Liu A, Plank L, Booth M. Differential Changes in Gut Microbiota After Gastric Bypass and Sleeve Gastrectomy Bariatric Surgery Vary According to Diabetes Remission. Obes Surg. 2017;27(4):917–925. doi:10.1007/s11695-016-2399-2

36. Wadden T, Berkowitz R, Womble L, Sarwer D. New England Journal CREST. N Engl J Med. 2005;353(20):2111–2120. doi:10.1056/NEJMoa050156

37. Van Gaal L, Dirinck E. Pharmacological approaches in the treatment and maintenance of weight loss. Diabetes Care. 2016;39(August):S260–S267. doi:10.2337/dcS15-3016

38. Jakicic JM, Kitabchi AE, Knowler WC. The long-term effectiveness of a lifestyle intervention in severely obese individuals. Am J Med. 2013;126(3):236–242. doi:10.1016/j.amjmed.2012.10.010.

39. Saxon DR, Iwamoto SJ, Mettenbrink CJ, et al. Antiobesity Medication Use in 2.2 Million Adults Across Eight Large Health Care Organizations: 2009-2015. Obesity. 2019;27(12):1975–1981. doi:10.1002/oby.22581

40. Rohde U, Hedbäck N, Gluud LL, Vilsbøll T, Knop FK. Effect of the EndoBarrier Gastrointestinal Liner on obesity and type 2 diabetes: A systematic review and meta-analysis. Diabetes, Obes Metab. 2016;18(3):300–305. doi:10.1111/dom.12603

41. De Jonge C, Rensen SS, Verdam FJ, et al. Endoscopic duodenal-jejunal bypass liner rapidly improves type 2 diabetes. Obes Surg. 2013;23(9):1354–1360. doi:10.1007/s11695-013-0921-3

42. Escalona A, Pimentel F, Sharp A, et al. Weight loss and metabolic improvement in morbidly obese subjects implanted for 1 year with an endoscopic duodenal-jejunal bypass liner. Ann Surg. 2012;255(6):1080–1085. doi:10.1097/SLA.0b013e31825498c4

43. Glaysher MA, Mohanaruban A, Prechtl CG, et al. A randomised controlled trial of a duodenal-jejunal bypass sleeve device (EndoBarrier) compared with standard medical therapy for the management of obese subjects with type 2 diabetes mellitus. BMJ Open. 2017;7(11). doi:10.1136/bmjopen-2017-018598

44. Betzel B, Drenth JPH, Siersema PD. Adverse Events of the Duodenal-Jejunal Bypass Liner: a Systematic Review. Obes Surg. 2018;28(11):3669–3677. doi:10.1007/s11695-018-3441-3

